# A missense variant in *PAOX* in American Staffordshire Terriers with juvenile-onset polyneuropathy

**DOI:** 10.1101/2025.10.16.682890

**Authors:** Lucie Chevallier, Garrett Bullock, Thibaut Troupel, Nicolas Blanchard-Gutton, Inès Barthélémy, Kaspar Matiasek, Frank Steffen, Ambre Courtin, Liz Hansen, Matthias Christen, Vidhya Jagannathan, Caroline Dufaure de Citres, Marion Signoret, Catherine Escriou, Marie Abitbol, Tosso Leeb, Dennis P O’Brien, Stéphane Blot, Laurent Tiret, Martin L Katz

## Abstract

Hereditary polyneuropathies in dogs mirror many features of human Charcot-Marie-Tooth (CMT) disease and offer valuable spontaneous models for discovery of genes’ variants involved in this disorder. A juvenile-onset polyneuropathy (JOP) in American Staffordshire Terriers (ASTs) resembles CMT, with motor and sensory deficits, often accompanied by laryngeal paralysis. To identify the underlying genetic cause, we conducted a genome-wide association study (GWAS) with 24 affected and 53 control ASTs that identified a significant locus at the distal end of chromosome 28. Homozygosity mapping analysis further defined a critical interval spanning 11 protein-coding genes and whole-genome sequencing revealed a missense variant in the *PAOX* gene encoding polyamine oxidase (NC_049249.1:g.41474541G>A), leading to a glycine-to-arginine substitution at a highly conserved residue, XP_038435135.1:p.(Gly382Arg). Genotypes at this variant were homozygous alternate in 96% of the affected dogs (n=53) and either homozygous reference or heterozygous in 336 unaffected ASTs; the mutant allele was not detected in any of 2519 genomes of other dog breeds. In silico predictions consistently supported its pathogenicity, and structural modeling suggested altered substrate binding. Although *PAOX* expression was preserved, proteomic profiling in affected nerve tissue revealed reduced levels of myelin-associated proteins, as well as dysregulation of mitochondrial and proteasomal pathways. Our findings establish this *PAOX* missense variant as the likely cause of JOP in ASTs and highlight the breed as a valuable large-animal model for studying inherited peripheral neuropathies. These results expand the spectrum of genes implicated in polyneuropathies and provide a basis for genetic testing and informed breeding strategies in ASTs.

**Author summary:** We studied a hereditary peripheral nerve disorder in American Staffordshire Terriers characterized by exercise intolerance, coordination problems, general weakness and difficulty breathing due to laryngeal paralysis. These signs closely resemble a group of inherited diseases in people known as Charcot-Marie-Tooth disease. To understand the genetic basis of this condition in dogs, we analyzed the genomes of affected and healthy individuals. Our research led us to a single gene, *PAOX*, which carries a missense variant likely affecting the function of polyamine oxidase and ultimately disrupting how the nerve cells function and survive. Most affected dogs had two copies of the mutant allele, while it was absent or in only one copy in healthy dogs. Additional analyses suggested that the variant may interfere with the nerves’ ability to maintain their structure and function properly. The mutant allele does not appear to reduce the overall level of PAOX protein, but it may affect how the protein works. Our findings provide a new genetic test that can help breeders avoid producing affected puppies. They also highlight this condition in dogs as a valuable model for understanding similar diseases in humans.

## Introduction

Charcot-Marie-Tooth disease (CMT), a group of hereditary motor and sensory polyneuropathies, is the most prevalent category of inherited peripheral neuropathies in humans. Despite significant genetic heterogeneity—over 100 causative loci have been identified (1)—CMT typically manifests as a slowly progressive, length-dependent sensorimotor polyneuropathy (PN). The disease commonly follows an autosomal dominant inheritance pattern, although X-linked and autosomal recessive forms also exist. Clinical subtypes are distinguished by age of onset, mode of inheritance, and severity of symptoms (2–5).

Comparable hereditary PNs have been documented in at least 22 dog breeds, often with juvenile or young-adult onset and slowly progressive clinical courses (6). These canine PNs frequently include motor or sensory deficits. The autonomic nervous system seems to be less frequently involved. Laryngeal paralysis is a frequently reported clinical sign in dogs with inherited peripheral neuropathy and may sometimes be the initial clinical manifestation of the disease (7–13). Laryngeal paralysis reflects the involvement of the recurrent laryngeal nerve and causes multiple clinical signs with respiratory dysfunction (stridor), exercise intolerance and gagging as the most frequent (14,15). Surgical intervention via arytenoid laryngoplasty (“tie-back”) is often effective in improving respiratory function and quality of life (11,16,17). Neurotization comprises the only alternative neurosurgical approach, albeit rarely used in practice. Syndromic neuropathies resembling Warburg micro syndrome have been described in the Alaskan Husky, Black Russian Terrier and Rottweiler, linked to variants in the *RAB3GAP1* gene (18–21). Nonsyndromic forms resembling human CMT (22,23) have been reported in the Dalmatian, Leonberger, Greyhound, Alaskan Malamute, Pyrenean Mountain Dog, Miniature Schnauzer and Labrador Retrievers among others (16,24–29). Causative variants have been identified in *NDRG1*, *SBF2 (MTMR13)*, *ARHGEF10*, *GJA9*, and *CNTNAP1*, several of which are also associated with human CMT subtypes (11,26,29–32). Notably, *MTMR2*, *MPZ*, and *SH3TC2* variants—genes known to cause human CMT—have been identified in Golden Retrievers with congenital hypomyelinating polyneuropathy (33). This genetic overlap underscores the translational relevance of canine models for understanding human inherited PNs.

A juvenile-onset form of polyneuropathy associated with laryngeal paralysis in American Staffordshire Terriers (ASTs) was described by Vandenberghe et al. (9). In this study, 14 affected dogs exhibited generalized motor and sensory deficits, with laryngeal paralysis observed in ten cases and megaesophagus in one case. Age at onset ranged from 1 to 6 months in most dogs, and the disease course was slowly progressive. Despite neuromuscular weakness, affected dogs could maintain a good quality of life, particularly when surgical treatment for laryngeal paralysis was performed. Electrodiagnostic testing was consistent with generalized, predominantly axonal and demyelinating, motor and sensory PN. Nerve histology was abnormal in all cases with myelinated fiber loss of variable severity. A notable histopathological feature seen in several cases was the presence of an inflammatory infiltrate, suggesting an immune-mediated component. Pedigree analysis supported an autosomal recessive mode of inheritance (9). The phenotype observed in ASTs shared many clinical and pathological features with human CMT, including juvenile onset, slow progression, and relatively preserved life expectancy (34), making it a valuable model for studying inherited peripheral neuropathies. In this study, we investigated the genetic basis of juvenile-onset polyneuropathy (JOP) in ASTs using a combination of genome-wide association analysis, homozygosity mapping, and whole-genome sequencing.

## Results

### Phenotypic confirmation and genetic screening

Dogs were classified in this study as affected or unaffected based on the presence or absence of the characteristic signs of JOP described previously (9). To determine whether the AST disorder was the result of a genetic variant previously associated with polyneuropathy in dogs, four affected ASTs (cases AST017, AST023, AST025, and AST026) were screened for previously reported variants associated with canine polyneuropathies. Specifically, they were tested for variants in the *RAB3GAP1*, *NDRG1*, *ARHGEF10*, and *GJA9* genes (20,21,26,29–31) and were found to be clear of the mutant alleles at all loci. These four dogs were also homozygous for the reference allele of the *ARSG* gene variant previously associated with cerebellar ataxia in ASTs (35). In addition, these four dogs, along with 15 other affected ASTs were genotyped for the *RAPGEF6* variant linked to laryngeal paralysis in Standard and Miniature Bull Terriers (detailed list in S1 Table). All tested individuals were homozygous for the reference allele at this locus (36).

### JOP inheritance pattern and pedigree analysis

Among the affected dogs recruited for this study, 16 were female and 13 were male in the French cohort; 13 were female and 9 were male (and one dog had no sex data available) in the U.S. cohort, and one was a male affected dog from Germany (see S1 Table). The sex distribution among the 52 affected dogs (with known sex data) included 23 males and 29 females. Under an autosomal mode of inheritance, an equal distribution of males and females (26 each) would be expected. A chi-square test showed no significant deviation from this expectation (p=0.41), indicating that the observed sex ratio is consistent with an autosomal pattern of inheritance. Pedigree information was obtained from owners when available. Analysis of 15 affected dogs’ pedigrees from the French cohort enabled the construction of a genealogical tree (see S1 Figure). As previously described by Vandenberghe et al. (2018)(9), the data confirmed that affected individuals were born to clinically unaffected parents. The pedigree also highlighted that inbreeding is a common practice among breeders. Notably, all 15 affected dogs shared a single common male ancestor born in the late 1970s and nominated best in breed within his first years of life. The presence of both affected males and females born to unaffected parents, together with the identification of inbreeding loops in multiple lineages, strongly supported an autosomal recessive mode of inheritance for this condition.

### Mapping of the JOP locus

#### Genome-Wide Association Study targets the end of CFA28

A GWAS was performed using 24 cases and 53 AST controls from the French cohort, and 117,409 markers were retained for analysis after quality filters. A significant association was identified at the distal end of chromosome 28, with two markers exceeding the Bonferroni-corrected genome-wide significance threshold (*P*_Bonf._ = 4.25 x 10^−7^). The two significantly associated SNVs in this GWAS were on CFA28 at 41,299,138 bp and 41,201,614 bp (CanFam4 assembly), with a likelihood ratio test *p*-value of 9.61 x 10^−9^ and 2.53 x 10^−7^ respectively. A Q-Q plot of expected and observed chi-squared values indicated that population stratification was successfully controlled (λ = 1.09) (Fig 1).

**Fig 1.**
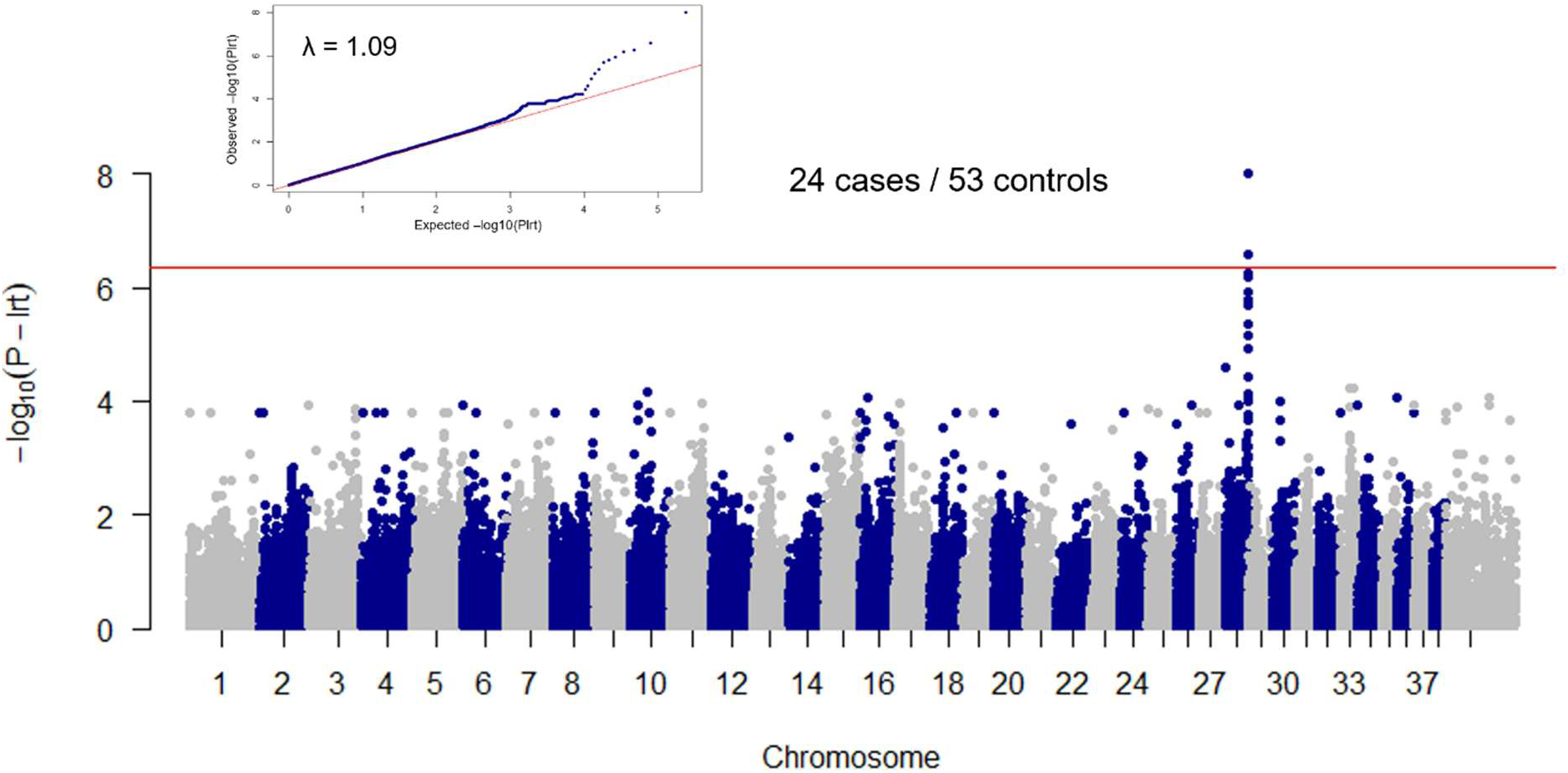
Results of the genome-wide association study. A GWAS was performed in a cohort of 24 JOP cases and 53 control dogs. The red line indicates the Bonferroni-corrected genome-wide significance threshold (*P*_Bonf_ = 4.25 x 10^-7^). The Manhattan plot shows a significant signal at the end of chromosome 28. The quantile-quantile (QQ) plot in the inset shows the observed (Y axis) versus expected (X axis) -log(*P*_lrt_) values. The straight red line in the QQ plot indicates the distribution of *P*_lrt_ values under the null hypothesis. The deviation of *p*-values on the right side indicates that these markers are more strongly associated with the trait than would be expected by chance. The genomic inflation factor (λ) was 1.09.

#### Linkage disequilibrium and homozygosity mapping define a critical interval for the JOP locus

Linkage disequilibrium (LD) analysis using the most significantly associated SNV (chr28:41,299,138) identified 18 neighboring markers in moderate to strong LD (r² ≥ 0.4), delineating a 1.42 Mb block (chr28:40,091,638–41,513,405) (Fig 2A, 2B). To further refine this locus, we extended the interval to the chromosome end and performed homozygosity mapping based on genotypes from 24 affected and 53 control dogs. The control group of unaffected ASTs was subdivided into 39 dogs of unknown genotype, referred to as “general controls,” and 14 dogs that had produced at least one affected offspring, referred to as “obligate carrier controls” (OCC). The critical interval was defined as the region between the last heterozygous marker preceding the shared homozygous stretch and the first heterozygous marker following it. Using this approach, we identified a 231,478 bp homozygous segment (chr28:41,362,645– 41,594,123) that was shared by 19 of 22 affected dogs with informative haplotypes; two affected dogs had missing data, and three carried different haplotypes (Fig 2B). This shared haplotype represents the critical interval for the JOP locus and encompasses 11 annotated protein-coding genes and six predicted genes according to NCBI release 106 (Fig 2C; see S2 Table for details).

**Fig 2.**
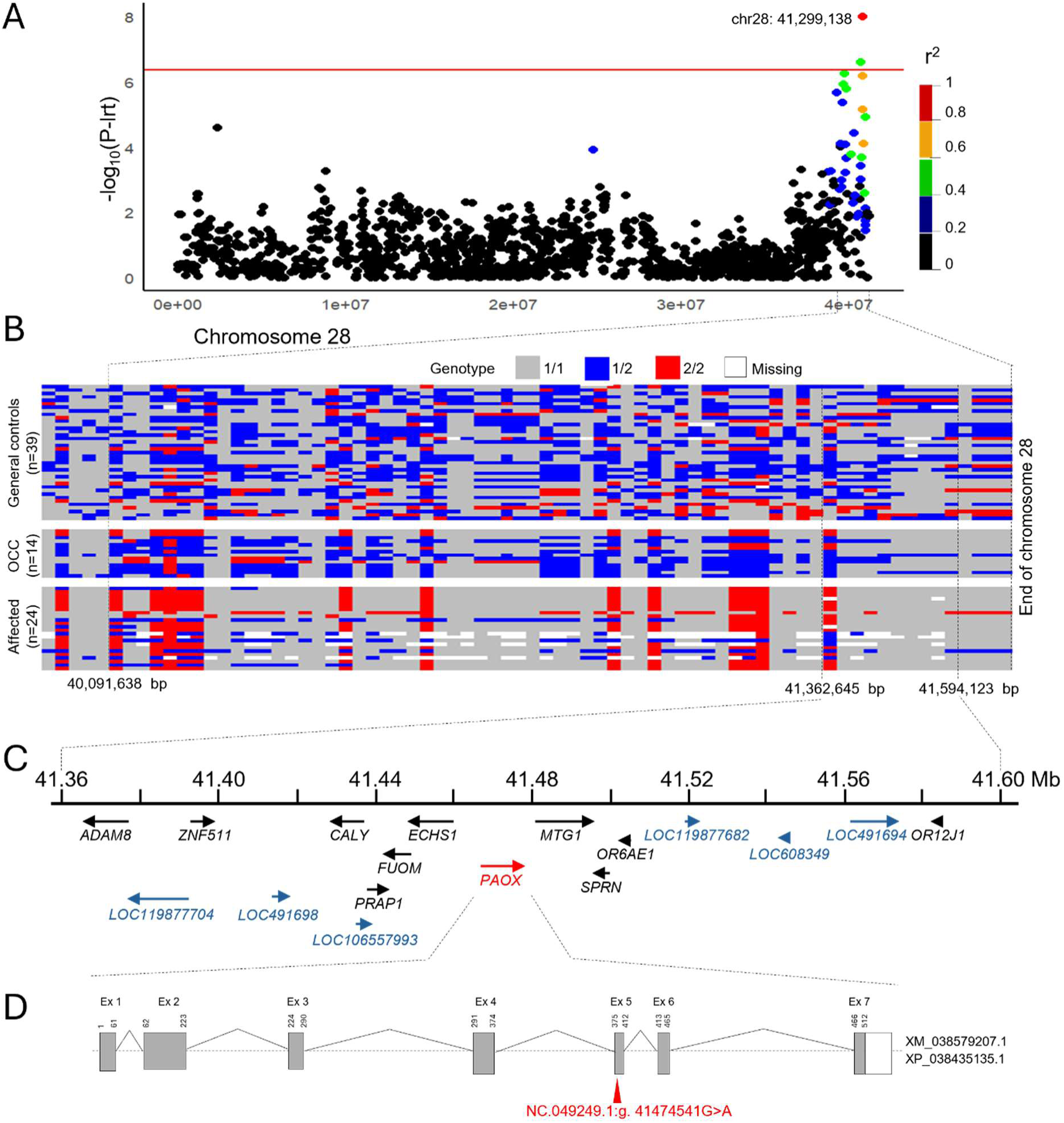
Mapping the JOP locus. (A) Linkage disequilibrium plot. Colors indicate the amount of linkage disequilibrium (LD) between the most associated marker (chr28: 41,299,138 in CanFam4 coordinates) and all the other markers spanning CFA28, ranging from red (*r*^2^ > 0.8) to black (*r*^2^ < 0.2). The interval of interest was determined based on LD values ≥ 0.4. (B) Heatmap depicting genotypic variation across SNVs on chromosome 28 for 77 dogs. The x-axis represents the SNV position in Mb, while the y-axis represents the dogs, grouped into three categories: general controls (n=39), obligate carrier controls (OCC; n=14), and affected (n=24). Each tile is color-coded according to the genotype of the respective dog at each SNV: homozygous for allele 1 (1/1) in gray, heterozygous (1/2) in blue, homozygous for allele 2 (2/2) in red, and missing genotypes in white. For each SNV, allele 1 was defined as the most frequent allele in the “general controls” group, and allele 2 as the less frequent allele in this group. (C) Gene annotation for the critical interval. The NCBI annotation release 106 listed 11 known protein-coding genes (in black or red) and 6 computer-predicted protein-coding genes (blue) in the critical interval. (D) Schematic representation of the *PAOX* gene showing the variant (NC.049249.1:g.41474541G>A) location in exon 5 (red arrow).

### Identification of candidate variants

Whole-genome sequencing (WGS) was performed on 10 ASTs affected with JOP, with an average coverage of 35.6X. Variant calling was conducted identifying 77,288,430 single nucleotide variants (SNVs) and small insertions/deletions (indels) across the 10 affected dogs prior to filtering.

To improve variant calling accuracy, the affected dogs were jointly genotyped with 323 non-AST dogs not affected with JOP. Variants were subsequently filtered to identify rare, coding-impactful candidates. Allele frequencies were calculated using a broader panel of 2,524 genomes from dogs of various breeds, including five unaffected ASTs (Table S2).

The variant filtering steps and the number of retained variants at each step are summarized in Table 1. First, variants failing standard quality control (QC) metrics were removed. Only variants in coding regions that passed QC were retained. Next, variants with low quality of sequencing depth were excluded. The remaining variants were filtered for predicted functional impact, retaining those classified as missense, nonsense, frameshift, splice-site, or otherwise protein-altering. Given that polyneuropathy is a rare phenotype but has been reported across multiple dog breeds (6), a minor allele frequency (MAF) cutoff of 0.05 was applied to focus on rare variants. The final filtering step retained only variants that were homozygous for the alternative allele in at least 8 of the 10 affected ASTs. This threshold was chosen to account for potential phenocopies, which are known to occur in canine PN (11,29,31). This filtering strategy identified a single missense variant in *PAOX* as a strong candidate (NC_049249.1:g.41474541G>A; XM_038579207.1:c.1144G>A; XP_038435135.1:p.(Gly382Arg)).

**Table 1.** Summary of variant filtering steps from whole-genome sequencing of ten JOP-affected ASTs.

| Filtering steps | Number of variants |
| --- | --- |
| Total variants (entire genome, 10 affected ASTs) | 77,288,430 |
| Standard QC filter, coding regions only | 41,482 |
| QD<sup>a</sup> ≥ 5; nonsynonymous coding variants | 31,589 |
| Allele frequency ≤ 0.05<sup>b</sup> | 4,664 |
| Homozygous in ≥ 8 of 10 samples | 1 |

<sup>a</sup> Quality by Depth (QD) score
<sup>b</sup> Based on allele frequencies across 2,524 genomes (including 5 unaffected ASTs).

Structural variants (SVs) were also investigated within the critical interval by visually inspecting short-read alignments of the 10 affected dogs using Integrative Genomics Viewer (IGV). No SVs unique to JOP- affected dogs were detected in this region. All BAM files from dogs used in this study are available (see Table S2).

### Genotype-phenotype association of the PAOX variant

We confirmed that all ten JOP-affected dogs subjected to whole-genome sequencing were homozygous for the mutant *PAOX* allele, using PCR and Sanger sequencing. To further assess the segregation and frequency of this allele, we genotyped 1,963 ASTs (389 with detailed phenotype records and 1,574 without). A total of 389 ASTs, comprising 269 dogs from the French cohort (29 JOP cases, 223 general controls, and 17 obligate carriers controls—defined as healthy parents of affected offspring), 111 dogs from the US cohort (23 cases and 88 general controls), and 9 dogs from the Swiss cohort (1 JOP case and 8 general controls), each with detailed phenotypic records were genotyped (see S1 Table). In both cohorts, all general control dogs carried either the G/G (wild-type) or G/A (heterozygous) genotype, and all obligate carrier controls were heterozygous (G/A), consistent with Mendelian recessive inheritance. Among the 53 JOP-affected dogs, 51 (96%) were homozygous for the variant allele (A/A), while 2 (4%) were G/G, indicating that the *PAOX* variant accounts for the majority, but not all, of the JOP cases (Table 2). Comparison of genotype frequencies between JOP cases and controls revealed a highly significant enrichment of the A/A genotype in affected dogs under a recessive model (US cohort: p = 2.71 × 10⁻^24^; French cohort: p = 4.25 × 10⁻³^5^), supporting a strong association between the *PAOX* variant and JOP in ASTs.

**Table 2.** Genotype frequencies for the *PAOX* variant (NC_049249.1:g.41474541G>A) in control and JOP-affected ASTs.

|  | JOP status | G/G<br>(ref/ref) | G/A<br>(ref/mut) | A/A<br>(mut/mut) | Total<br>(n=) |
| --- | --- | --- | --- | --- | --- |
| US cohort | Affected | 0% (0) | 0% (0) | 100% (23) | 23 |
|  | General control <sup>a,b</sup> | 81.8% (72) | 18.2% (16) | 0% (0) | 88 |
| French cohort | Affected | 6.9% (2) | 0% (0) | 93.1% (27) | 29 |
|  | General control <sup>a,b</sup> | 91.9% (205) | 8.1% (18) | 0% (0) | 223 |
|  | Obligate carrier control <sup>a,c</sup> | 0% (0) | 100% (17) | 0% (0) | 17 |
| Swiss cohort | Affected | 0% (0) | 0% (0) | 100% (1) | 1 |
|  | General control <sup>a,b</sup> | 100% (8) | 0% (0) | 0% (0) | 8 |
| Global population tested at MU <sup>d</sup> | Unknown phenotype | 81.9% (525) | 11.5% (74) | 6.6% (42) | 641 |
| Global population tested at Antagene <sup>d</sup> | Unknown phenotype | 86.4% (806) | 11.6% (108) | 2% (19) | 933 |

The number of dogs is indicated in brackets.
<sup>a</sup> ASTs $\geq 3$ years old that showed no signs of PN.
<sup>b</sup> Controls with no descendant information were designated as “general controls”.
<sup>c</sup> Control dogs were classified as “obligate carrier controls” if they were healthy parents of affected offspring.
<sup>d</sup> ASTs without known phenotype status that were submitted for various purposes to MU (US) or for genetic testing of the JOP locus to Antagene (France).

In addition, we analyzed two additional cohorts of ASTs with unknown phenotypes. The first cohort comprised 641 dogs submitted to the University of Missouri (MU) for reasons unrelated to this study. Among these, 42 were homozygous mutant (A/A), 74 heterozygous (G/A), and 525 homozygous reference (G/G), resulting in a carrier frequency of 12.4% (74/599) and a mutant allele frequency of 12.3% (158/1282). Although this group represents a broader sample of U.S. ASTs, it may not reflect the general U.S. population, as the MU archive does not constitute a random sample of the breed. The second cohort included 933 ASTs submitted to Antagene for genetic screening of the *PAOX* variant. Among these, 19 were A/A, 108 G/A, and 806 G/G, corresponding to a carrier frequency of 11.8% (108/914) and a mutant allele frequency of 7.8% (146/1866). While this group provides an overview of the European AST population, it may not represent the general European population due to non-random sampling of dogs submitted to Antagene (Table 2).

### Conservation and predicted impact of the PAOX variant

Because the NC_049249.1:g.41474541G>A variant in the *PAOX* gene was the only candidate variant left after filtering and further showed a strong genotype–phenotype association, we focused on this variant. This SNV identified in exon 5 of the *PAOX* gene encoding polyamine oxidase (Fig 2D) was predicted to result in a glycine-to-arginine substitution at amino acid position 382 of the PAOX protein, XP_038435135.1:p.(Gly382Arg). The affected residue showed high sequence conservation across vertebrates (S2 Figure). Glycine is conserved at this position in nearly all species, except in rats, which carry an alanine residue—a small, neutral substitution only marginally larger than glycine. A conservation score of 9.236 was obtained using PhyloP, based on a comparison of 100 vertebrate species (37).

The PAOX:p.Gly382Arg variant was predicted to be deleterious by four different in silico pathogenicity prediction tools (Table 3). According to SIFT, the amino acid substitution is classified as damaging if the score is ≤ 0.05 (38). For the other tools—MutPred2, PredictSNP, and PolyPhen-2—a score approaching 1 suggests a higher probability that the variant is deleterious. (39–41).

**Table 3.** Predicted impact of PAOX missense variant. . Output of different *in silico* protein prediction tools for the PAOX:XP_038435135.1:p.G382R missense variant.

| Prediction tool | Variant score / Score for deleterious prediction | Prediction of pathogenicity |
| --- | --- | --- |
| PredictSNP | 86 % | Deleterious* |
| MutPred2 | 0.816† | Likely pathogenic |
| SIFT | 0 | Deleterious |
| PolyPhen-2 | 0.991‡ | Probably damaging |

\* Each of the individual programs in the PredictSNP suite classifies the p.G382R missense variant as deleterious.
† False positive rate of 5% at this cutoff.
‡ Sensitivity of 0.5 and specificity of 0.95.

In humans, the orthologous residue is Gly381. Pathogenicity prediction using the AlphaMissense tool (42) assigned a score of 0.967 to the Gly381Arg substitution in human PAOX, which exceeds the threshold of 0.564 for likely pathogenicity. Three-dimensional structural modelling of the wildtype and mutant canine PAOX proteins indicated that the p.Gly382Arg substitution introduces a bulky and positively charged residue close to the putative binding site of the positively charged polyamine substrates. This might lead to an electrostatic and/or steric repulsion of the substrate (S2 Figure).

### PAOX expression analysis in peripheral nerves of affected and control dogs

#### Western blot analysis

To assess whether PAOX protein expression was altered in peripheral nerves of affected dogs, western blot analysis of total proteins was conducted using an anti-PAOX antibody directed against an N-terminal epitope. A ∼60 kDa band was detected in both control and affected dogs. Band intensities appeared similar between affected and control samples across all nerve types (Fig 3, upper panel). Total protein staining was used to confirm equal protein loading (Fig 3, lower panel). Densitometric analysis showed that, after normalization to total protein, PAOX band intensities in affected dogs were 102%, 99%, and 99% of control levels in the sciatic, fibular, and ulnar nerves, respectively. Therefore, we concluded that the variant has no impact on PAOX protein abundance in nerves.

**Fig 3.**
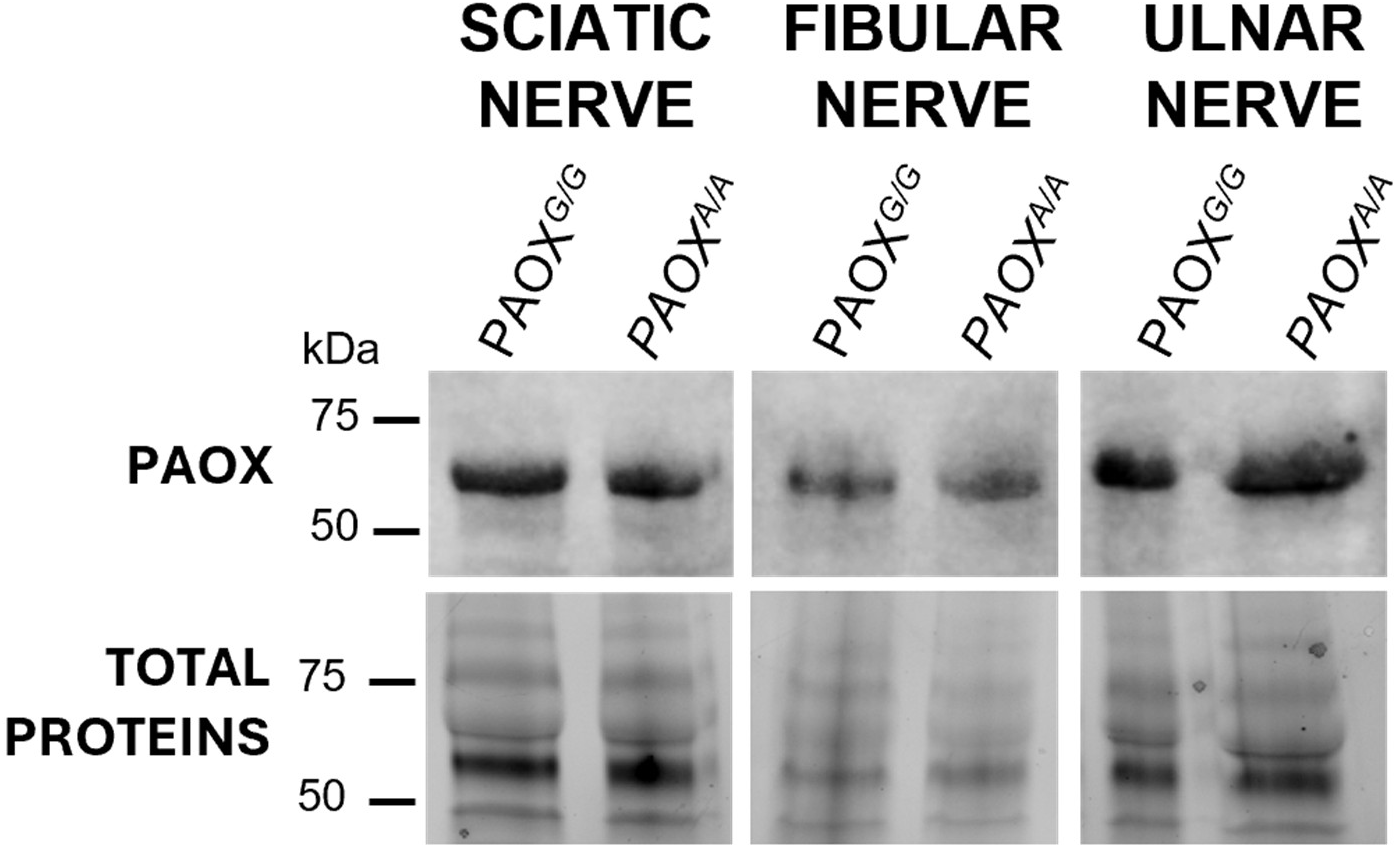
PAOX protein expression in nerves of control (G/G) and affected (A/A) dogs. Western blot analysis of sciatic, fibular, and ulnar nerve extracts using an N-terminal anti-PAOX antibody shows similar expression of PAOX protein (∼60 kDa) in control Golden Retrievers (GR1 for sciatic and fibular nerves and GR2 for ulnar nerve: *PAOX^G/G^*) and affected ASTs (AST007 for sciatic and fibular nerves and AST009 for ulnar nerves: *PAOX^A/A^*). The lower panel shows similar staining of proteins between 40 and 80 kDa.

### Fluorescence immunohistochemistry analysis

To further assess PAOX protein spatial distribution, we co-immunostained peripheral nerve transverse sections of one affected AST (AST011) and one control Golden Retriever (GR1) with antibodies raised against PAOX and neurofilament 68 (NF-L, a marker of axonal structure) (43,44) (Fig 4). Consistent with western blot results, PAOX immunoreactivity in affected nerves could be detected at levels comparable to the control. The overall spatial staining patterns of PAOX and NF-L were similar between affected and control dogs.

**Fig 4.**
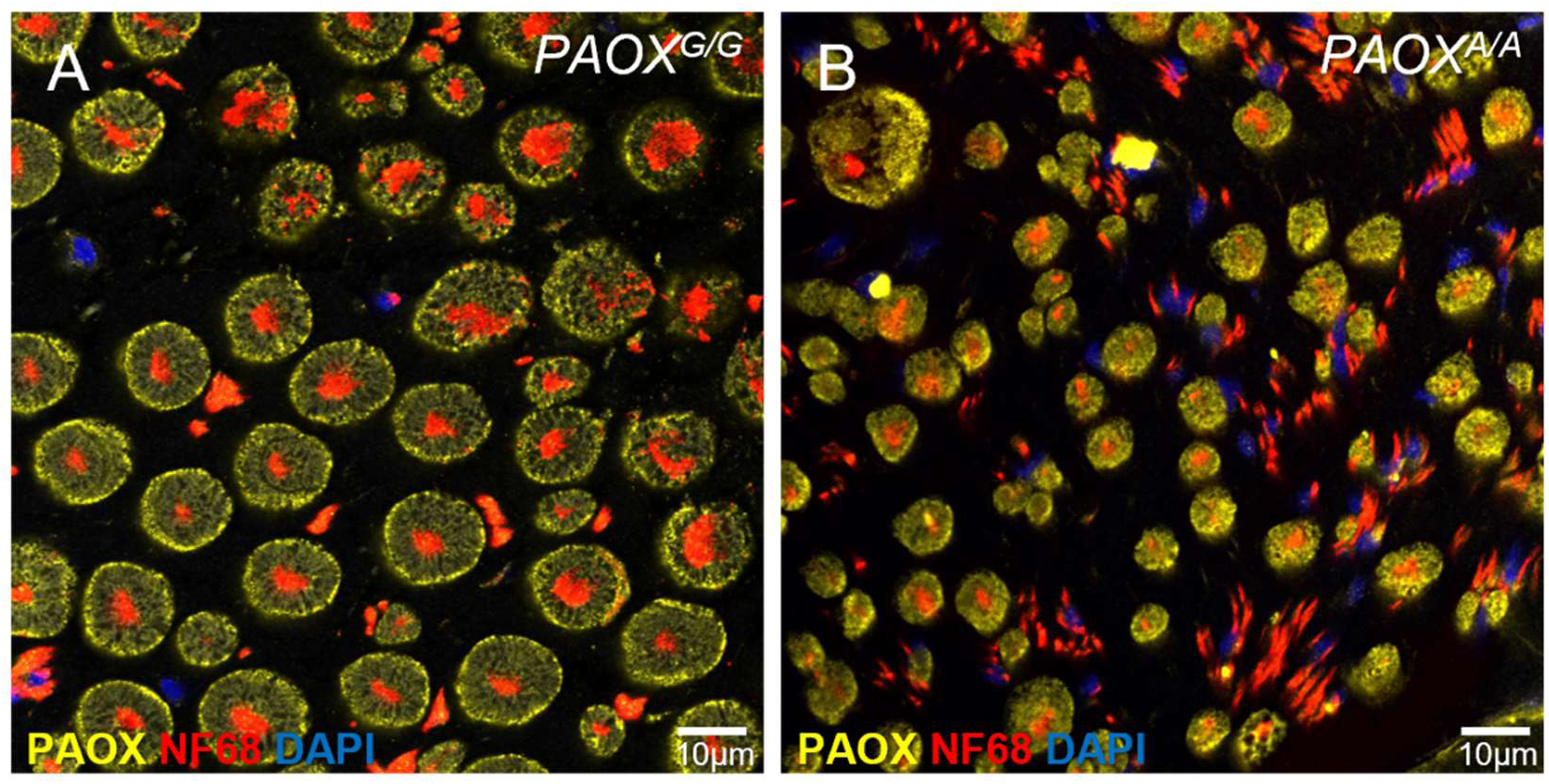
Immunolabelling of PAOX and NF68 in fibular nerve sections. Fluorescence micrographs showing PAOX expression (yellow), NF68 (red), and DAPI nuclear staining (blue) in a control dog (*PAOX^G/G^*) (A) and an affected AST (*PAOX^A/A^*) (B). Scale bar 10 μm.

In control nerves, PAOX localized predominantly to Schwann cells (Fig 4A). In affected dog, because of extensive nerve degeneration, nerve fibers appeared reduced in diameter, although NF-L and relative PAOX staining intensity was conserved. Notably, isolated NF-L staining was more prominent in affected nerves, suggesting a higher proportion of solitary axons. DAPI staining indicated increased cellularity in affected dog, consistent with the cell infiltrate previously described by Vandenberghe et al. (9). Overall, fluorescence histochemistry revealed unchanged PAOX expression but features of axonal loss, Schwann cell loss, and increased cellular infiltration in affected dog (Fig 4B).

### Protein profiling in affected and control nerves

The nerves’ total protein profiles obtained in electrophoresis between affected and control dogs were globally similar except for a marked reduction of a major protein band between 25 and 37 kDa in the affected dogs, a difference consistently observed in the sciatic, fibular, and ulnar nerves electrophoretic profiles (Fig 5A). To further investigate this difference, the corresponding band from sciatic nerve samples from one *PAOX* homozygous mutant affected AST (AST007; MS2) and one Golden Retriever control (GR1; MS1) was excised and analyzed by label-free quantitative mass spectrometry. A total of 239 proteins were identified in at least one sample (MS1 or MS2). Fold changes were calculated as the log₂ ratio of MS2 to MS1 peak areas (S3 Table). Of these, 143 proteins showed substantial differences in abundance between MS1 and MS2 (|log₂FC| > 1), with 87 proteins downregulated in the affected nerve. Gene Ontology (GO) enrichment analysis using PANTHER revealed significant (FDR < 0.05) overrepresentation of processes related to mitochondrial fission and peripheral nervous system myelination, including axon ensheathment, Schwann cell development, and peripheral nerve system development. Key proteins included *PLEC* (plectin), *PRX* (periaxin), *COL6A1* (collagen type VI alpha 1 chain), *THY1* (Thy-1 cell surface antigen), *CDK5* (cyclin dependent kinase 5), *MPZ* (myelin protein zero), *MBP* (myelin basic protein), and *LOC487015* (myelin P2 protein). The most abundant protein in MS1, based on spectral count, was myelin protein P0 (XP_545771.1), which was more than six times as abundant in MS1 compared to MS2, with a >29-fold difference based on peptide peak area (|log₂FC| = 4.89). DAVID functional clustering also highlighted mitochondrion and myelin sheath as top categories, along with cytoskeleton, lipid metabolism, and neuron apoptotic processes.

**Fig 5.**
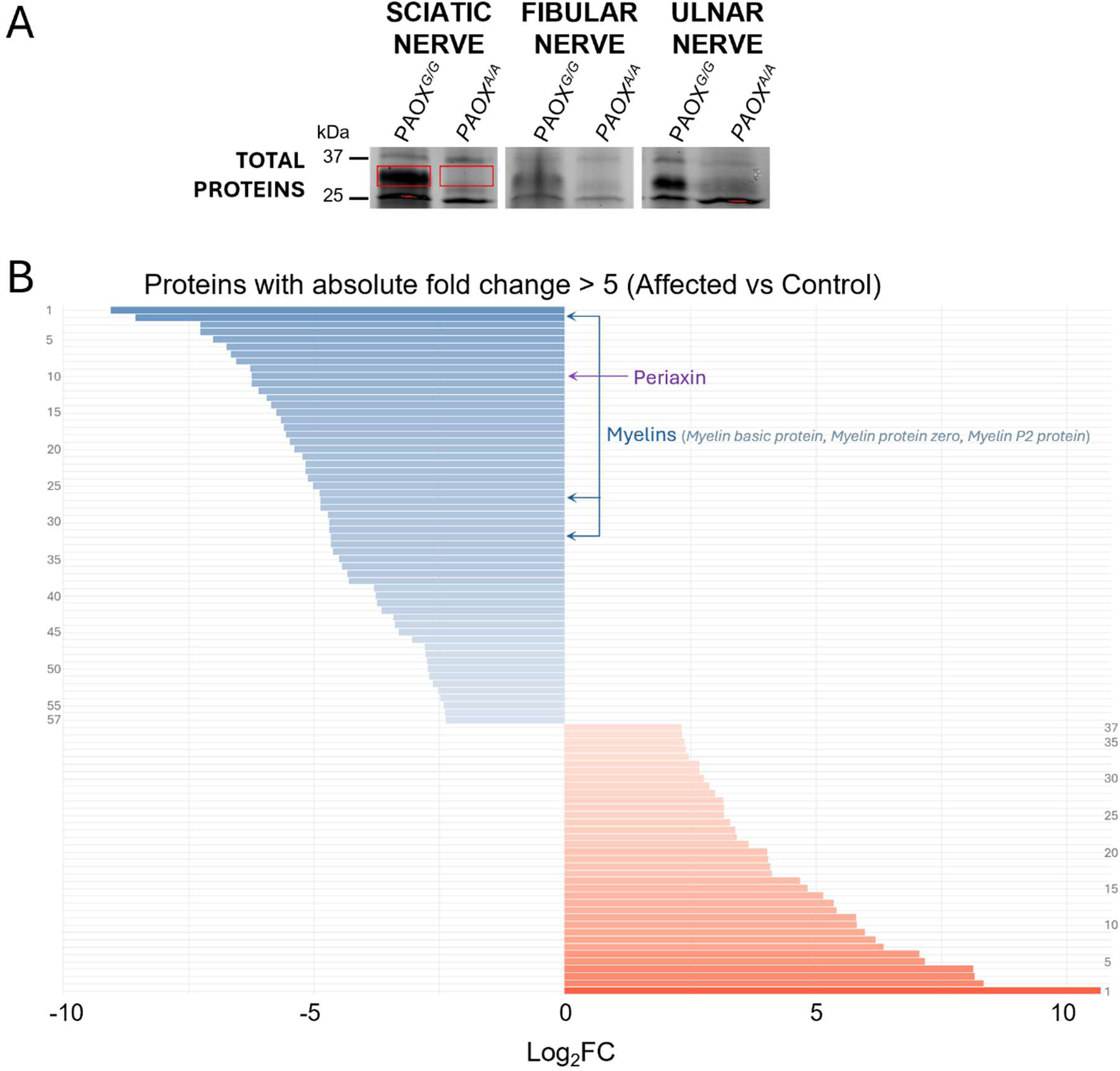
Differential protein abundance in the sciatic nerve of an affected (*PAOX^A/A^*) and a control dog (*PAOX^G/G^*). (A) Representative image of protein electrophoresis gel showing a distinct band of differential intensity between 25 and 37 kDa, suggesting altered protein expression in a JOP-affected dog compared to a control dog in sciatic, fibular and ulnar nerves. (B) Bar plot of proteins with an absolute fold change > 5 between affected and control samples, based on label-free quantification from mass spectrometry (log₂ fold change > 2.32). Proteins are ordered by log₂ fold change, with positive values indicating increased abundance in the affected dog and negative values indicating decreased abundance. The numbers on the left and right correspond to the down- and up-regulated proteins, respectively, listed in S3 Table.

In contrast, 56 proteins were upregulated in the affected nerve (MS2). While GO enrichment for this subset did not reach statistical significance, the top nominally enriched terms included phospholipid efflux, transport regulation, and proteasomal protein catabolic processes. Several proteasome subunits were upregulated (e.g., *PSMA3*, *PSMA5*, *PSMA6*, *PSMA7*, *PSME2*, *PSMD8*). DAVID analysis identified the proteasome and neurodegeneration pathways as the most enriched.

The most differentially expressed proteins (fold change > 5) are listed in Fig 5B. We concluded that the *PAOX* variant has no visible impact on PAOX abundance and localization but was associated with an extensive nerve remodeling and degeneration with significant myelin loss and altered regulatory pathways.

## Discussion

Our combined GWAS and WGS approach in three independent American Staffordshire Terrier (AST) cohorts identified a missense variant in *PAOX* (NC_049249.1:g.41474541G>A) whose segregation explains approximately 96% of juvenile-onset polyneuropathy (JOP) cases in this breed. The GWAS mapped the trait to the distal region of CFA28 containing 11 protein-coding genes including *PAOX*, while genome-wide WGS filtering in affected dogs independently identified a single compelling candidate variant in *PAOX*. The convergence of these two independent approaches, the absence of other plausible pathogenic variants, and the predicted deleterious functional impact of the missense change together support *PAOX* as the major locus for JOP in ASTs. Nevertheless, around 4% of diagnosed dogs do not carry the mutated *PAOX* allele in a homozygous state, indicating that the phenotype is genetically heterogeneous with less frequent pathogenic variants that remain to be further identified. Genetic heterogeneity is well-documented in canine peripheral neuropathies, both across and within breeds (6). In Leonberger dogs with polyneuropathy, three characterized variants account for only ∼50% of cases (11,29,31). Similarly, Cook et al. (33) reported locus heterogeneity within a single breed caused by variants in different genes. The availability of a *PAOX* gene test now enables systematic collection of JOP cases homozygous for the reference allele, which may increase the size of a cohort that will facilitate the discovery of additional causative genes in ASTs.

PAOX is a flavin adenine dinucleotide (FAD)-dependent amine oxidase that catalyzes the oxidation of N1- acetylated polyamines, preferentially N1-acetylspermine over N1-acetylspermidine, generating spermidine or putrescine, 3-acetoamidopropanal, and hydrogen peroxide (45,46). This catalytic activity has been well characterized biochemically, yet its precise physiological role—particularly within the nervous system—remains incompletely understood, and no in vivo animal model, nor human phenotype, with impaired PAOX activity has been described. Our finding that the PAOX variant is associated with peripheral neuropathy suggests that the protein may also have other direct or indirect functions that are specific to neural axons or Schwann cells.

The canine variant identified here is predicted to result in a p.Gly382Arg substitution adjacent to Phe383 (S2 Figure). Structural studies of murine PAOX (47) indicate that the homologous residue Phe375 lies within a hydrophobic subpocket that accommodates N1-acetylspermine. Replacing glycine with a bulky, positively charged arginine in this region is predicted to disrupt the hydrophobic environment, creating electrostatic and steric repulsion with the positively charged polyamine substrates. This structural change may impair substrate binding and catalytic efficiency. Conservation of Gly382 across vertebrate species (S2 Figure B), combined with the high AlphaMissense pathogenicity score of 0.967 for the homologous human Gly381Arg substitution (42) and other computational pathogenicity predictions, strongly supports the interpretation of this change as deleterious. Additional population data from gnomAD v4.1.0 (48) reinforces this conclusion: a Gly381Glu substitution—introducing a negatively charged glutamic acid—has been reported at an allele frequency of only 1.67 × 10⁻⁵ (27 of 1,612,606 alleles) and never in the homozygous state. Other rare missense variants affecting neighboring residues were also identified exclusively in the heterozygous state and at extremely low frequencies. The ClinVar database (49,50) lists over 70 SNVs in human *PAOX*; however, no pathogenicity data are reported for any of them, and none involve codon 381. These observations collectively suggest that amino acid changes in this region of PAOX are poorly tolerated and likely to be pathogenic. While these predictions are compelling, direct measurement of PAOX enzymatic activity in affected dogs will be essential to confirm the biochemical consequences of the substitution.

Protein analysis provides further support for a pathogenic link between the PAOX variant and peripheral nerve degeneration. Western blotting with an anti-PAOX antibody revealed no obvious difference between affected and control dogs in the abundance of the PAOX protein, but total protein profiling uncovered the absence in affected dogs of a prominent 25–37 kDa band present in controls. Mass spectrometry identified this band as containing myelin protein zero (MPZ), neurofilament heavy and medium polypeptides, tubulin β, and periaxin—proteins critical for maintaining axonal stability and myelin integrity. MPZ variants are well-established causes of CMT1B and CMT2 in humans (51–56) and have been implicated in canine polyneuropathy (33), and periaxin variants have been linked to CMT4F in several families (57–60). The depletion of these structural proteins in affected ASTs aligns closely with the histopathological evidence of myelin loss and axonal degeneration (9), suggesting that disruption of polyamine homeostasis in *PAOX*-mutant dogs may indirectly destabilize axonal and myelin components.

The biology of PAOX must be viewed in the broader context of polyamine metabolism. Polyamines— putrescine, spermidine, and spermine—are small, positively charged molecules with essential roles in DNA stabilization, chromatin remodeling, mRNA translation, ion channel modulation, oxidative stress control, and immune regulation (46,61–65). Their intracellular levels are tightly controlled through a balance between biosynthesis, catabolism, and transport (46,63). In catabolism, spermidine/spermine N1- acetyltransferase (SAT1) acetylates spermine and spermidine, generating substrates for PAOX and spermine oxidase (SMOX) (45). Disruption of this pathway is increasingly recognized as a contributor to neurodegeneration. Zahedi et al. (2020) (66) demonstrated that double knockout of *Smox* and *Sat1* in mice causes progressive ataxia, Purkinje cell loss, cerebellar demyelination, and neuroinflammation, with profound alterations in polyamine levels in both cerebellum and cerebrum (66). Polyamine imbalance has also been linked to Parkinson’s disease (67,68), Alzheimer’s disease (69–72), Snyder-Robinson syndrome (73), Bachmann-Bupp syndrome (74), epilepsy (75,76), and traumatic brain injury (77), through mechanisms that include oxidative stress from hydrogen peroxide and aldehyde by-products, ion channel dysfunction, and altered DNA conformation (78–80). Moreover, polyamine metabolism, specifically involving the molecule spermine, was found to be abnormally elevated in the neuropathic forms of mucopolysaccharidoses. High levels of spermine were identified in the cerebrospinal fluid of both animal models and human patients with neuropathic mucopolysaccharidoses subtypes but not in non-neuropathic forms (81). These findings reinforce the concept that balanced polyamine turnover is essential for neuronal survival, myelin preservation and homeostasis of the nervous system.

Polyamines are also important modulators of immune functions, capable of suppressing pro-inflammatory cytokine production (82,83) and influencing macrophage polarization (84). Human families with SAT1 loss of function variants showed typical features of systemic lupus erythematosus, such as increased type I interferon signaling, kidney inflammation, and autoantibody production (85). In *Smox/Sat1* knockout mice, neurodegeneration is accompanied by CD4+ and CD8+ T cell infiltration in the leptomeninges (66), suggesting that altered polyamine catabolism can trigger inflammatory responses in the nervous system. This is particularly relevant to AST JOP, where inflammatory infiltrates are observed in many cases as a consequential overlap neuropathy (9)—a feature not infrequently seen in other breed-associated canine polyneuropathies and consistent with the immunomodulatory roles of polyamines.

The identification of a *PAOX* variant as a major cause of JOP in ASTs has important comparative and translational implications. Given the extensive genetic heterogeneity of Charcot–Marie–Tooth disease (CMT) in humans (1), *PAOX* should now be considered a candidate gene in undiagnosed CMT patients. The AST JOP phenotype offers a large-animal model of mixed axonal and demyelinating peripheral neuropathy, creating opportunities to investigate the role of polyamine metabolism in nerve biology and to evaluate potential therapeutic strategies—ranging from dietary modulation to small-molecule inhibitors or supplementation (46,64,65,86).

In conclusion, our study establishes, for the first time, a link between *PAOX* and polyneuropathy in vertebrates. Affected ASTs constitute a valuable spontaneous model of CMT that can be leveraged for mechanistic studies and preclinical therapeutic trials (87). These findings expand the repertoire of genes associated with inherited neuropathies and highlight the critical importance of polyamine metabolism in maintaining peripheral nerve structure and function.

## Materials and Methods

### Dog phenotyping, pedigrees analysis and sample collection

Phenotyping and sample collection were conducted at multiple institutions: the University of Missouri (MU; Columbia, MO, USA), the National Veterinary School of Alfort (École Nationale Vétérinaire d’Alfort, ENVA; Maisons-Alfort, France), the Centre Hospitalier Vétérinaire Frégis (CHVF; Paris, France), VetAgro Sup (VAS; Marcy l’Étoile, France), Antagene (La Tour-de-Salvagny, France) and the Vetsuisse Faculty of the University of Zurich (Zurich, Switzerland). A total of 53 American Staffordshire Terriers (ASTs) with juvenile-onset polyneuropathy (JOP) were included in this study. Of these, 23 were phenotyped at MU, 29 at French institutions (ENVA, CHVF, VAS) and one in Switzerland. Among the French cohort, 12 of the 14 dogs originally reported by Vandenberghe et al. (2018) (9) were included. Additional cases were examined at ENVA (n=5), CHVF (n=3), and VAS (n=1). Phenotyping was performed following the same diagnostic protocol described by Vandenberghe et al. (2018) (9), including neurological examination, electromyography (EMG), and nerve biopsy. Eight other affected dogs were diagnosed by primary care veterinarians based on clinical signs and laryngoscopy and were included via ENVA or Antagene. Control dogs were ASTs aged ≥3 years with no signs of neurological disease. Dogs confirmed to be clinically healthy but that had produced at least one affected offspring were designated as “obligate carrier controls”; other controls with no descendant information were designated as “general control”. Recruited dogs were either purebred ASTs or AST-like, based on morphology. Sex ratios were compared to the expected 1:1 distribution under an autosomal mode of inheritance using a chi-square test in R Statistical Software (v. 4.3.2). Pedigree data were collected when available. Familial relationships were analyzed, and genealogical trees were constructed using GenoPro 2020 (version 3.1.0.1). For clarity, dogs phenotyped or sampled in the United States are referred to as the “US cohort” (n=111), those collected in France are referred to as the “French cohort” (n=269) and those collected in Switzerland are termed the “Swiss cohort” (n = 9).

### DNA and nerve tissue collection

DNA samples were collected from affected and control dogs across all cohorts, with informed owner’s consent. For the French and Swiss cohorts, genomic DNA was isolated from EDTA-anticoagulated blood or buccal swabs using the Maxwell® 16 Blood or Buccal Swab LEV DNA Purification Kits (Promega, Dübendorf, Switzerland). For the US cohort, DNA was extracted from EDTA blood using protocols previously described (Katz et al., 2005). DNA quantity and quality were assessed by NanoDrop ND1000 spectrophotometry (NanoDrop ND1000, Thermo Scientific) and samples were stored at ≤ –20 °C.

Nerve tissue was collected from three JOP-affected dogs (AST007, AST009, AST011; listing in S1 Table 1) evaluated at ENVA who were euthanized at owner request by pentobarbital overdose (Dolethal, Vetoquinol, France) due to poor quality of life. Due to owner restrictions, full necropsy was not performed; collected tissues included sciatic and fibular nerves (AST007), fibular nerve (AST011), and ulnar nerve (AST009). Control nerve samples (sciatic, fibular, and ulnar) were obtained post-mortem from two healthy Golden Retrievers (GR1 and GR2) euthanized for unrelated reasons. All nerve samples were snap-frozen in isopentane pre-cooled in liquid nitrogen and stored at –80 °C for downstream analysis.

### Genotyping of known variants in PN-associated genes

Genetic tests LPN1 (c.1955_1958+6del10bp in *ARHGEF10* gene,), LPN2 (c.1107_1108delAG in *GJA9* gene), AMPN (c.293G>T in *NDRG1* gene), NDRG1 (c.1080_1089del10bp in *NDRG1* gene), JLPP (c.743delC in *RAB3GAP1* gene) and NCL-A (c.296G>A, *ARSG* gene) (20,21,26,29–31,35) were performed by Antagene laboratory (Antagene, La Tour-de-Salvagny, France). The *RAPGEF6* gene variant (c.1793_1794ins36) associated with laryngeal paralysis in Bull Terriers was genotyped in 19 affected ASTs (see S1 Table) as described in Hadji Rasouliha et al 2019 (36).

### Genome-wide Association Study (GWAS)

55 ASTs (24 cases and 31 controls) from the French cohort (see S1 Table for a detailed listing of the dogs) were genotyped with the CanineHD BeadChip array (Illumina Inc) containing 220,853 markers. 22 control ASTs that had been genotyped for a former project using the CanineHD BeadChip (Illumina Inc) containing 174,376 markers were included (35). As these markers were originally derived from the CanFam2.0 assembly, their genomic positions were converted to CanFam3.1. Data were merged and pruned using the program PLINK (v1.07), (88). 77 individuals and 154,140 common markers were kept for analysis (raw genotypes are available for download at https://doi.org/10.5281/zenodo.17370812). Quality control (QC) on genotyping data was performed. Samples with missingness > 10% were removed, and only SNVs with a minor allele frequency greater than 5% and a genotyping rate greater than 90% were included. SNV genomic positions were converted from CanFam3.1 to CanFam4 using the Liftover tool from UCSC genome browser (genome-euro.ucsc.edu). After QC and conversion, the final dataset included a total of 77 dogs (24 cases and 53 controls) and 117,409 SNV markers. GWAS was performed using a univariate mixed model with a standardized relatedness matrix in GEMMA v.0.97 (89), and likelihood ratio test *p*- values (P-lrt) were used to determine significantly associated markers. A Bonferroni-corrected significance threshold (α = 0.05) was calculated using all SNVs included in the analysis. Genomic inflation factor, as a measure of controlling for population genetic structure, was calculated in R Statistical Software (v. 4.3.2). Linkage disequilibrium (LD) between the top associated SNV in the GWAS and other SNVs on chromosome 28 was calculated using Plink (v1.07), (88). Manhattan, Q-Q, and linkage disequilibrium (LD) plots, as well as the processing and visualization of genotyping data, were generated in R (v. 4.3.2) using the qqman, dplyr, and ggplot2 packages (90). All genomic positions are reported with respect to the dog reference genome assembly CanFam4 and NCBI annotation release 106 unless otherwise specified.

### Short-read whole-genome sequencing

Ten cases (see S1 Table for a detailed list of dogs) were whole-genome sequenced. DNA samples were submitted to the University of Missouri Genomics Technology Core Facility for library preparation and 2 x 150 base pair paired-end sequencing on an Illumina Nova-Seq 6000 instrument with an average target coverage depth of 35.6X. Raw sequence reads were aligned to the Dog10K_Boxer_Tasha (GCA_000002285.4, Ensembl Release 107) reference genome assembly with the Burrows-Wheeler Aligner (BWA). The resulting Sequence Alignment Map (SAM) files were converted to Binary Alignment/Map (BAM) files and sorted using SAMtools (v. 1.11). PCR duplicates were marked with PicardTools (v. 2.23.8) and a modified GATK (v. 3.8) best practice pipeline was used for realignment and recalibration of .bam files and for var and to generate genome variant call format (gVCF) files for each sample in conjunction with whole-genome sequence data obtained from 323 dogs of other breeds that had not exhibited signs of polyneuropathy (S4 Table). Variants in each sample were called individually with GATK HaplotypeCaller in gVCF mode and joined with GATK CombineGVCFs. Joint genotyping was performed using GATK GenotypeGVCFs, and functional predictions of the called variants were annotated with SnpEff (91). SnpSift software (92) was used to filter out low quality variants from the WGS dataset and to extract variants from the affected samples. Extracted variants were tabulated with GATK VariantsToTable to generate a browsable variant report relative to the reference sequence. Nonsynonymous variants (missense, nonsense, and potentially other coding-impactful types) located in coding regions with a Quality by Depth (QD) score ≥ 5 were retained for downstream analysis. Variants with an allele frequency (AF) ≤ 0.05 were retained based on 2524 genomes (internal controls and publicly available genomes, 5 ASTs included, see S4 Table). Candidate variant positions were then converted to Canfam4. Furthermore, the critical interval was analyzed for structural variants by visual inspection of the alignment files of JOP cases compared to controls using Integrative Genomics Viewer (IGV) (93).

### In silico analysis of variant and protein predictions

PredictSNP, MutPred2, SIFT and PolyPhen-2 in silico prediction tools were used to predict biological consequences of the discovered missense variant NC.049249.1:g.41474541G>A leading to amino acid substitution PAOX:XP_038435135.1:p.G382R on the encoded protein (38–41). Alphamissense pathogenicity tool (42) was used to evaluate the predicted pathogenicity of a homologous variant PAOX protein (PAOX:Gly381Arg) in the human (Q6QHF9 code in Uniprot). The Genome Aggregation Database (gnomAD, v4.1.0) (48) was searched for the corresponding variant in the human PAOX gene (NP_690875.1). The protein models were created with the ColabFold program (94). Nucleotide conservation across different vertebrate species was evaluated using Multiz Alignment and PhyloP from the PHAST package on the UCSC Genome Browser (95). Predicted protein molecular weight based on amino acid sequence was calculated using the Protein Molecular Weight tool from bioinformatics.org (www.bioinformatics.org). All references to the canine *PAOX* gene correspond to the accessions NC_049249.1 (NCBI accession), XM_038579207.1 (mRNA), and XP_038435135.1 (protein) unless otherwise specified.

### Targeted Genotyping

A custom TaqMan SNV genotyping assay was designed to confirm the presence of the candidate single base substitution in *PAOX* (NC.049249.1:g.41474541G>A). PCR primer sequences for the assay were *5’*- TCTCCCCACACAGGTCTGT-*3’* and *5’*-TCGGTCAGAGTCTCCATGAACT-*3’* and competing probe sequences were *5’*-*VIC*-TCCTCTGCGGGTTCA-*NFQ-3’* (reference allele) and *5’*-*FAM*-TCCTCTGCAGGTTCA-*NFQ*-*3’* (mutant allele). Assays were performed in 25 µL reaction mixtures consisting of the DNA sample, 72 µM primers, 16 µM probes in TaqMan Universal PCR Master Mix (Thermo Fisher Scientific, Waltham, MA, USA) on a StepOnePlus Real-Time PCR System (Applied Biosystems, Thermo Fisher Scientific, Waltham, MA, USA).

Genotype frequencies between JOP cases and controls were compared separately in the French and US cohorts using Fisher’s exact test, implemented in R (v. 4.3.2). Analyses were conducted under a recessive genetic model, in which dogs homozygous for the mutant allele (mut/mut) were classified as “affected genotype,” while those with either the reference homozygous (ref/ref) or heterozygous (ref/mut) genotypes were grouped as “non-affected genotype”. The null hypothesis tested was that the proportion of mut/mut genotypes is equal between cases and controls.

### SDS-gel electrophoresis and western blotting

Protein extracts (25 μL) were loaded on a 4-15% SDS PAGE stain free gel (BioRad, Hercules, CA, USA). After gel activation, proteins were transferred to PVDF membrane with iBlot2 dry blotting system (Invitrogen by Thermo Fisher Scientific, Waltham, MA, USA). Membrane was incubated with a rabbit anti-PAOX antibody (SAB2105175; 1:1000; Sigma-Aldrich, St Louis, MO, USA), as well as with peroxidase-coupled swine anti-rabbit immunoglobulins (P039901-2; 1:2000; Agilent, Santa Clara, CA, USA). The immunoblots were revealed with the SuperSignal^TM^ West Femto kit (Thermo Fisher Scientific, Waltham, MA, USA), and monitored on a BioRad Chemidoc Imager (BioRad, Hercules, CA, USA). To ensure equal loading of protein samples, total protein normalization was used as a loading control. After gel activation, total protein levels across all lanes were visualized using a BioRad Chemidoc Imager (BioRad, Hercules, CA, USA). Densitometry analysis was performed using ImageJ (Schneider 2012), and the total lane protein signal was used to normalize the intensity of target protein bands. This approach was chosen to avoid the variability in housekeeping proteins whose expression could vary due to nerve degeneration in the affected dogs (96). For each sample lane, the area under the curve (AUC) corresponding to the total protein signal was measured, as well as the AUC for the PAOX-specific band. The PAOX signal was then normalized to the total protein content by calculating the ratio of the PAOX band AUC to the total protein AUC for each lane. This normalization was performed separately for each of the three nerve types analyzed. For each nerve type, the normalized PAOX value of the control dog was set to 100%, and the corresponding value for the affected dog was expressed as a percentage of the control.

### Fluorescence immunohistochemistry

Cryosections of nerves (7 µm) were collected on Superfrost slides (Thermo Fisher Scientific, Waltham, MA, USA), air-dried, and fixed in a 1:1 acetone/methanol solution at -20°C. Sections were rehydrated in PBS for 10 min and blocked in 6% PBS/BSA for 30 minutes at room temperature. Slides were incubated overnight at 4°C with a primary anti-PAOX antibody solution (1:100, SAB2105175; Sigma-Aldrich, St. Louis, MO, USA or 1:100, STJ96463; St John’s Laboratory, London, UK) in 3% PBS/BSA. Following overnight incubation, the slides were washed in PBS and incubated for 1 hour at room temperature with a rhodamine-conjugated anti-rabbit secondary antibody (1:500, #111-026-047; Jackson ImmunoResearch Laboratories, Inc., West Grove, PA, USA). After additional washes, sections were incubated with a primary anti-NF68 antibody solution (1:200, NCL-NF68-DA2; Novocastra, Newcastle upon Tyne, UK) for 1 hour at room temperature and washed in PBS. The slides were then incubated in Alexa Fluor 647 anti-mouse secondary antibody solution (1:500, #ab150119; AbCam plc, Cambridge, UK) for 1 hour at room temperature and in PBS. Finally, sections were incubated with DAPI (#32670; Sigma-Aldrich, St. Louis, MO, USA) for 5 minutes, rinsed, and mounted using Fluoromount™ (Dako, Glostrup, Denmark). Images were acquired using a Leica STELLARIS 5 confocal fluorescence microscope (Leica Biosystems, Wetzlar, Germany). Image processing was performed using Fiji software (v. 2.16.0) (97).

### Proteomic analysis by liquid chromatography–tandem mass spectrometry

Protein extracts (25 μg) from sciatic nerve of one control (GR1) and one JOP-affected dog (AST007) were denatured (5 minutes, 95 °C), separated on a 4–15% SDS-PAGE gel (Bio-Rad), and stained with Coomassie Brilliant Blue. After destaining (9% ethanol, 9% acetic acid), a band between 25–37 kDa was excised and digested using in-gel Coomassie protocol (described here: https://docs.research.missouri.edu/proteomics/pro_in-gel_digestion_1d_coom.pdf). Peptides were extracted, desalted using C18 tips (Pierce Cat# 87774), lyophilized, and resuspended in 5 μL of 5% acetonitrile, 0.1% formic acid. One microliter was injected onto a PepSep15 C18 column (15 cm × 150 μm × 1.5 μm; Bruker) using a Bruker nanoElute system coupled to a timsTOF PRO with a CaptiveSpray source. Peptides were separated at 600 nL/min with a 30 minutes gradient: 3–17% B (0–10 min), 17–25% B (10–15 min), 25–37% B (15–20 min), 37–80% B (20–23 min), flushing (80–40% B) and re-equilibration (3% B, 26–30 min). MS data were acquired in positive-ion PASEF mode (TIMS on), over m/z 100–1700 (calibrated 03/24/2025), with 1 MS and 10 MS/MS frames per 1.17 s cycle. Target MS intensity was 10,000 counts/sec (min threshold 250); dynamic exclusion with re-acquisition if precursor increased >4× after 0.4 min; isolation width was 2 m/z (<700 m/z) or 3 m/z (800–1500 m/z); rolling collision energy ranged from 76– 123% of 42.0 eV. Raw .d files were searched using PEAKS Studio v12 against the *Canis familiaris* NCBI database (trypsin, max 2 missed cleavages, carbamidomethyl-Cys fixed, Met-Ox and N/Q-deamidation variable mods, 20 ppm precursor and 0.05 Da fragment tolerances). FDR was <1% at peptide and protein levels. To minimize carryover, three blank runs were inserted between samples. Protein abundance between the control dog (MS1) and the JOP-affected dog (MS2) was measured as peak area, and fold change was calculated as the log₂ ratio of MS2 to MS1 (log₂[Area_MS2 / Area_MS1]). To account for missing values, undetected protein intensities were imputed using half the minimum non-zero intensity observed across both samples. Proteins with an absolute log₂ fold change greater than 1 (i.e., fold change > 2 or < 0.5) were considered differentially abundant. Biological process enrichment analysis was performed using the Gene Ontology (GO) resource (https://geneontology.org/) (98), and functional annotation was conducted using DAVID Bioinformatics Resources (https://davidbioinformatics.nih.gov/) (99), focusing on proteins with fold changes greater than 2 or less than 0.5. A subset of proteins with a log₂ fold change greater than ∼2.32 (i.e., fold change > 5) was selected for visualization. Bar plot displaying log₂ fold change values was generated using the ggplot2 package in R (v4.3.2) (90).

## Acknowledgments

Authors are very grateful to owners and breeders who participated in the study. We thank the French Biological Resources Center Cani-DNA for the biobanking of ENVA AST samples. This study was supported by funding from CRB-Anim (PIA1 2012–2022, GIS CRB-Anim 2022–2027) and IBISA (2022–2024). We are grateful to Solène Diop and Manon Guehl for their assistance with nerve tissue sample collection, to Cindy Maenhoudt and Guillaume Robiteau for their assistance with sampling non affected dogs and to Andrea Carlier for her valuable support with western blotting and fluorescence immunohistochemistry imaging. We also thank Maud Rimbaud for her help with array data management. Finally, we wish to honor the memory of Gary S. Johnson, who sadly passed away during the course of this project, and whose contributions were essential to many aspects of this work.

## Accession IDs

Dog, *Canis lupus familiaris*, NCBI Taxon ID 9615.

*PAOX*, Peroxisomal N(1)-acetyl-spermine/spermidine oxidase [Canis lupus familiaris (dog)] gene ID 480798.

## Declarations

### Author contribution information

Conceptualization: LC, GB, MK

Data curation: LC, GB, GSJ, CDC, AC, MA, LH, VJ

Formal analysis: LC, GB, TL, NBG

Investigation: SB, TT, DPO, MS, CE, KM, IB, FS, MC

Resources: NBG

Writing – original draft: LC, GB, TL, MK Writing – review & editing: All authors Supervision: LC, MK

Funding acquisition: LC, LT, MK

### Ethics approval

All animal procedures complied with local regulations. Work conducted at the University of Missouri was approved by the Institutional Animal Care and Use Committee (IACUC). For the French cohort, IACUC approval was not required because all the dogs were clinical patients at four different veterinary hospitals and sampling was performed as part of their routine diagnostic procedures, as stipulated by French regulations. The sampling of healthy control dogs of the Swiss cohort was approved by the Canton of Bern (BE94/2022). All examinations were performed with the owners’ informed consent. Additional whole- genome sequencing data were obtained from publicly available sources or from internal datasets generated for other projects.

### Data availability

The SNV genotype data for 77 American Staffordshire Terriers (ASTs) generated in this study have been deposited in Zenodo and can be accessed via DOI: https://doi.org/10.5281/zenodo.17370812. Whole- genome sequencing files reported in this paper can be found in the NCBI Sequence Read Archive (SRA Bioproject no. PRJNA263947). The whole-genome sequences for ten American JOP-affected ASTs are available in SRA under accession numbers: SRS4394520, SRS5354887, SRS5354857, SRS11384554, SRS11384556, SRS11384557, SRS11384558, SRS11384574, SRS11384576 and SRS11384606 (see S4 Table for more details). The genome sequence of the case from the Swiss cohort is available under accession SAMEA4867927 at the European Nucleotide Archive.

## Supporting information titles and captions

**S1 Table. Phenotype and genotype information of all ASTs included in this study (French and US cohorts).**

**S1 Figure. Family tree of a subset of ASTs affected by JOP.**

**<u>Caption:</u> Family pedigree showing a subset of ASTs included in the study.** Squares represent males and circles represent females. Filled symbols indicate dogs affected by JOP; unfilled symbols indicate unaffected dogs or dogs without phenotype information. A common male ancestor shared by the affected individuals in this family is annotated in red. Not all affected dogs from the study are represented in this pedigree (Dog ID are listed in S1 Table).

**S2 Table. List of genes within the candidate region (chromosome28:41,362,645-41,594,123 bp, in CanFam4 coordinates).**

**S2 Figure. Details of the PAOX:p.Gly382Arg variant.**

**<u>Caption:</u>** (**A**) Multispecies alignment of PAOX amino acid sequences in the region of the variant illustrates relatively high sequence conservation across vertebrates. The alanine residue in rats is neutral and only marginally larger than the glycine seen in most species. Sequences represent accessions NP_690875.1 (*H. sapiens*), XP_038435135.1 (*C. familiaris*), NP_001013620.2 (*B. taurus*), NP_722478.2 (*M. musculus*), NP_001099781.1 (*R. norvegicus*), XP_003641516.1 (*G. gallus*) and XP_690593.2 (*D. rerio*). (**B**) Three-dimensional structure models of the wildtype (left) and mutant (right) canine PAOX protein. The upper half of the protein corresponds to the FAD binding domain, whereas the lower half represents the catalytic domain with the hydrophobic substrate binding pocket. The p.Gly382Arg substitution is immediately adjacent to ^383^Phe (highlighted in green). The homologous murine residue, ^375^Phe, has been shown to line a hydrophobic subpocket contacting the N^1^-acetylspermine substrate (Sjögren *et al.* 2017). The p.Gly382Arg substitution introduces a bulky and positively charged side chain, which may very well result in electrostatic and/or steric repulsion of the positively charged polyamine substrates of the enzyme. The protein models were created with the ColabFold program (Mirdita *et al.* 2022).

**S3 Table. Raw mass spectrometry data with log₂ fold change between affected and control nerves.**

**<u>Caption:</u>** Raw protein abundance data from mass spectrometry analysis of peripheral nerve tissue from a JOP-affected AST (MS2) and a healthy control dog (MS1). Protein abundance is expressed as peak area. Missing intensity values were imputed using half the minimum non-zero intensity across both samples. The log₂ fold change (log₂[MS2 / MS1]) was calculated using imputed values.

**S4 Table. Metadata for 2,534 whole-genome sequenced dogs.**

**<u>Caption:</u>** Metadata for 2,534 dogs from various breeds used as a reference panel for variant filtering. The table includes the BioSample ID, SRA accession, BioProject ID, breed, sex, Dog ID (listed in Table S1), and a “Remark” column indicating which dogs are JOP-affected. Among these, 16 ASTs are included, with 11 JOP-affected ASTs highlighted in red.

