## Supplementary figures and images for "A missense variant in *PAOX* in American Staffordshire Terriers with juvenile-onset polyneuropathy"

### S1 Figure

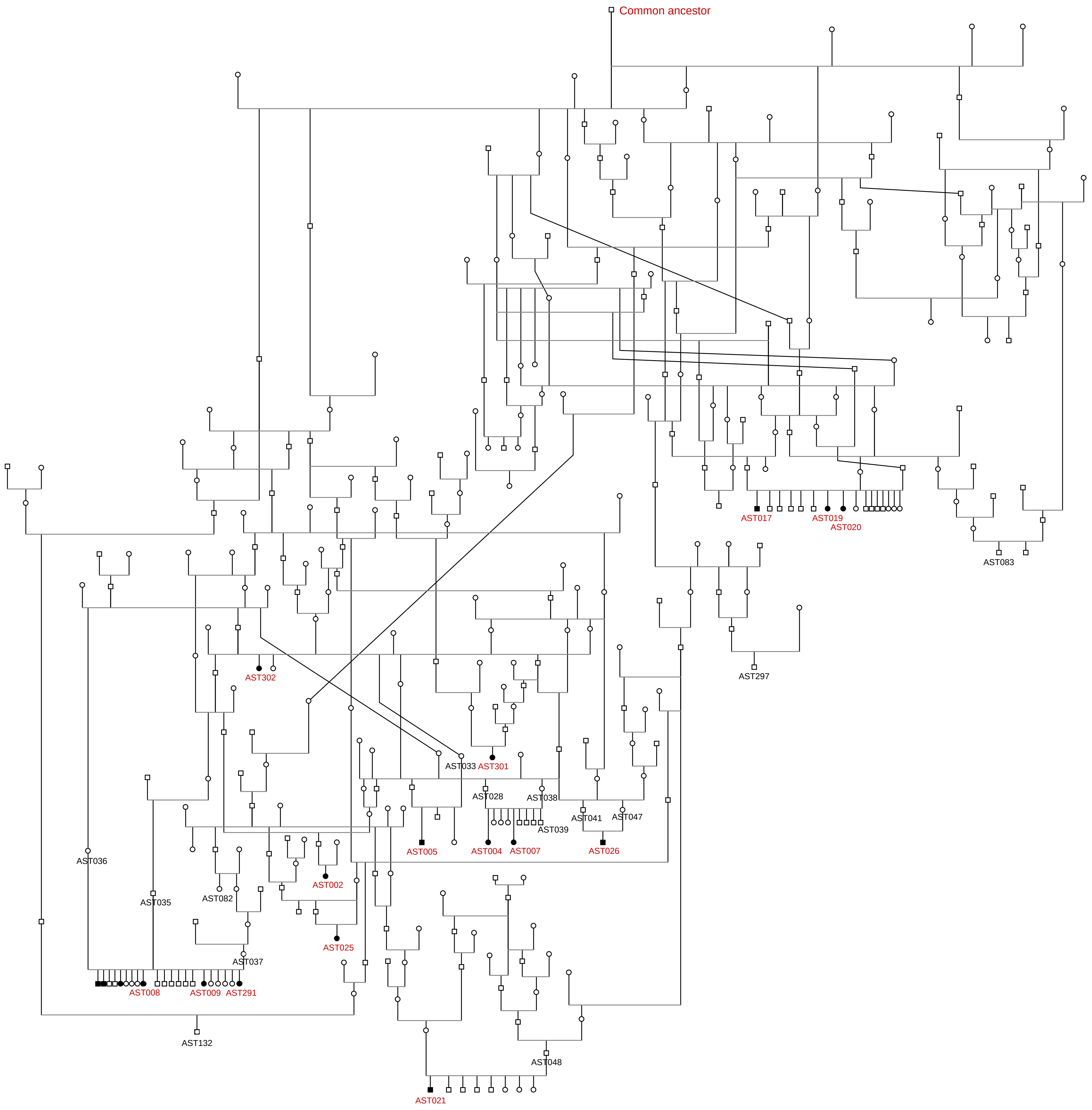
