## Supplementary material for "A missense variant in *PAOX* in American Staffordshire Terriers with juvenile-onset polyneuropathy": S2 Figure

**A**

```
                           p.Gly382Arg
                                ↓
H. sapiens      371 PAFASVHVLCGFIAGLESEFM 391
C. familiaris   372 .S.E................. 392
B. taurus       372 .P.QAS............... 392
M. musculus     364 .S.E.S............... 384
R. norvegicus   391 .S.E.S....A.....Q.... 401
G. gallus       358 .PEQLG..........K...Y. 378
D. rerio        370 .TERFG.....W...Q...Y. 390
```

**B**

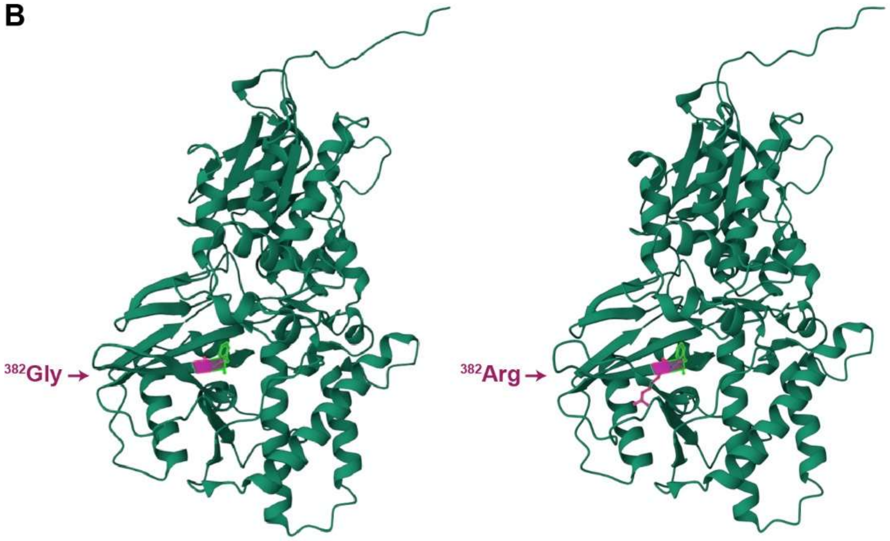


**Figure S2.** Details of the PAOX:p.Gly382Arg variant. (**A**) Multispecies alignment of PAOX amino acid sequences in the region of the variant illustrates relatively high sequence conservation across vertebrates. The alanine residue in rats is neutral and only marginally larger than the glycine seen in most species. Sequences represent accessions NP_690875.1 (*H. sapiens*), XP_038435135.1 (*C. familiaris*), NP_001013620.2 (*B. taurus*), NP_722478.2 (*M. musculus*), NP_001099781.1 (*R. norvegicus*), XP_003641516.1 (*G. gallus*) and XP_690593.2 (*D. rerio*). (**B**) Three-dimensional structure models of the wildtype (left) and mutant (right) canine PAOX protein. The upper half of the protein corresponds to the FAD binding domain, whereas the lower half represents the catalytic domain with the hydrophobic substrate binding pocket. The p.Gly382Arg substitution is immediately adjacent to $^{383}$Phe (highlighted in green). The homologous murine residue, $^{375}$Phe, has been shown to line a hydrophobic subpocket contacting the $N^1$-acetylspermine substrate (Sjögren *et al.* 2017). The p.Gly382Arg substitution introduces a bulky and positively charged side chain, which may very well result in electrostatic and/or steric repulsion of the positively charged polyamine substrates of the enzyme. The protein models were created with the ColabFold program (Mirdita *et al.* 2022).
